# Whisking asymmetry signals motor preparation and the behavioral state of mice

**DOI:** 10.1101/568030

**Authors:** Sina E. Dominiak, Mostafa A. Nashaat, Keisuke Sehara, Hatem Oraby, Matthew E. Larkum, Robert N.S. Sachdev

## Abstract

A central function of the brain is to plan, predict and imagine the effect of movement in a dynamically changing environment. Here we show that in mice head fixed in a plus-maze, floating on air, and trained to pick lanes based on visual stimuli, the asymmetric movement and position of whiskers on the two sides of the face signals whether the animal is moving, turning, expecting reward or licking. We show that 1) we can decode and predict the behavioral state of the animal based on this asymmetry, 2) that tactile input from whiskers indicates little about the behavioral state, and 3) that movement of the nose correlates with asymmetry, indicating that facial expression of the mouse is itself correlated with behavioral state. Amazingly, the movement of whiskers – a behavior that is not instructed or necessary in the task--informs an observer about what a mouse is doing in the maze. Thus, these mobile tactile sensors reflect a behavioral and movement-preparation state of the mouse.

## Introduction

One of the principal functions of the brain is to control movement (Llinas, 2001; Wolpert and Ghahramani, 2000; Wolpert and Landy, 2011). According to one view, brains may even have evolved for the sole purpose of guiding and predicting the effect of movement (Llinas, 2001). Whether or not the brain evolved for motor control, it is clear that the activity of many brain circuits is intimately linked to movement (Fetz, 1994) and that movement can involve many sensory-motor modalities. For example, the simple act of reaching to touch an object requires postural adjustments, and motion of the head, eye and limbs, with the eyes often moving first (Barnes et al., 1979; Anastasopoulos et al., 2009).

The rodent whisker system is a multimodal sensory-motor system. While it is often called a model sensory system (Van der loos and Woolsey, 1973), where each whisker is associated with 1000s of neurons in the trigeminal somatosensory pathways, this system is in fact also a model motor system – with single muscles associated with each whisker (Dorfl, 1983; Grinevich et al., 2005; Haidarliu et al. 2013). Not only do mice have the potential to control the motion of these tactile sensors individually, but the motion of the whiskers is often coordinated with motion of the head (Sachdev et al., 2002; Towal and Hartmann 2006; Grant et al., 2012; Mitchinson et al., 2011; Mitchinson and Prescott 2013; Schroeder and Ritt, 2016). Additionally, whisking can be triggered by sniffing, chewing, licking and walking (Welker 1964; Deschenes et al., 2012; Grant et al., 2012; Arkley et al., 2014; Sofroniew et al., 2014). Whisking can also be used to detect the location of objects, it can be used in social contexts, and it can predict direction of movement of the freely moving and head fixed animal (Krupa et al. 2004; Sellien et al. 2005; Knutsen et al. 2006; Godde et al. 2010; Cao, et al., 2012; Grant et al. 2012a,b; Sofroniew et a., 2014; Arkley et al. 2014; Reimer et al., 2014; Saraf-Sinik et al. 2015; Voigts et al. 2015; Lenschow et al., 2015). Taken together this earlier work, indicates that rodents move their whiskers in a variety of contexts; they move their whiskers while navigating through and exploring their environment; they move their whiskers during social interactions; and they move their whiskers when they change their facial expression.

The sensory motor circuits dedicated to whiskers are often studied in the context of active sensation, i.e. when the whiskers are moved to touch and detect objects or to discriminate between objects. Even though a lot is known about whisker use, most earlier studies in had fixed rodent have limited their observations to simple preparations. They have avoided studying whisker use in complex environments that have walls, contours and textures (but see Sofroniew et al., 2014). While the development of virtual reality systems has increased the complexity of behaviors used in head-fixed rodents, virtual systems are predominantly-geared toward creating virtual visual worlds around animals (Holscher et al. 2005; Harvey et al., 2009). Here we use an alternative platform that floats on air, one in which head fixed mice are trained to search for a dark lane. As they navigate the environment, they can touch, manipulate and experience it (Nashaat et al., 2016; Voigts et al., 2018). Here we began by looking for stereotypy in whisking during different behavioral epochs that occur in the course of the task. Once stereotypy became evident, we tested the obvious hypothesis that asymmetric whisking reflected tactile input from whiskers. Our work reveals that instead of a tactile and exploratory function during navigation in the maze, the asymmetric movement of whiskers, predicts the behavioral state of the animal.

## Methods

We performed all procedures in accordance with protocols approved by the Charité Universitätsmedizin Berlin and the Berlin Landesamt für Gesundheit und Soziales (LAGeSo) for the care and use of laboratory animals.

### Surgery

Five adult male mice, weighing 25-32g were used in these experiments. Animals were anesthetized with ketamine/xylazine (90 mg/kg ketamine and 10 mg/kg xylazine) to prepare them for head-fixation. A lightweight aluminum head post was attached using a mixture of Rely-X cement and Jet acrylic black cement. Animals were monitored during recovery and were given antibiotics (Enroflaxacin) and analgesics (Buprenorphine and Carprofen).

### Air Track Plus maze

The details of the custom-made plus maze, air-table and monitoring system for detecting the location and position of the plus maze have been published previously (Nashaat et al., 2016). Here we used a clear plexiglass air table mounted on aluminum legs, on which we placed a 3D printed circular platform 30 cm in diameter, that was shaped into a plus maze, where each lane of the maze was 10 cm long, 4 cm wide and 3 cm high, and the center of the maze, the “rotation area” was 10 cm in diameter (Figure 1A). The walls of the lanes had different textures. Some lanes had smooth walls others had walls with vertical evenly spaced raised indentations. Mice were not trained to discriminate between the different textures of the walls, they were simply exposed to them.

**Figure 1.**
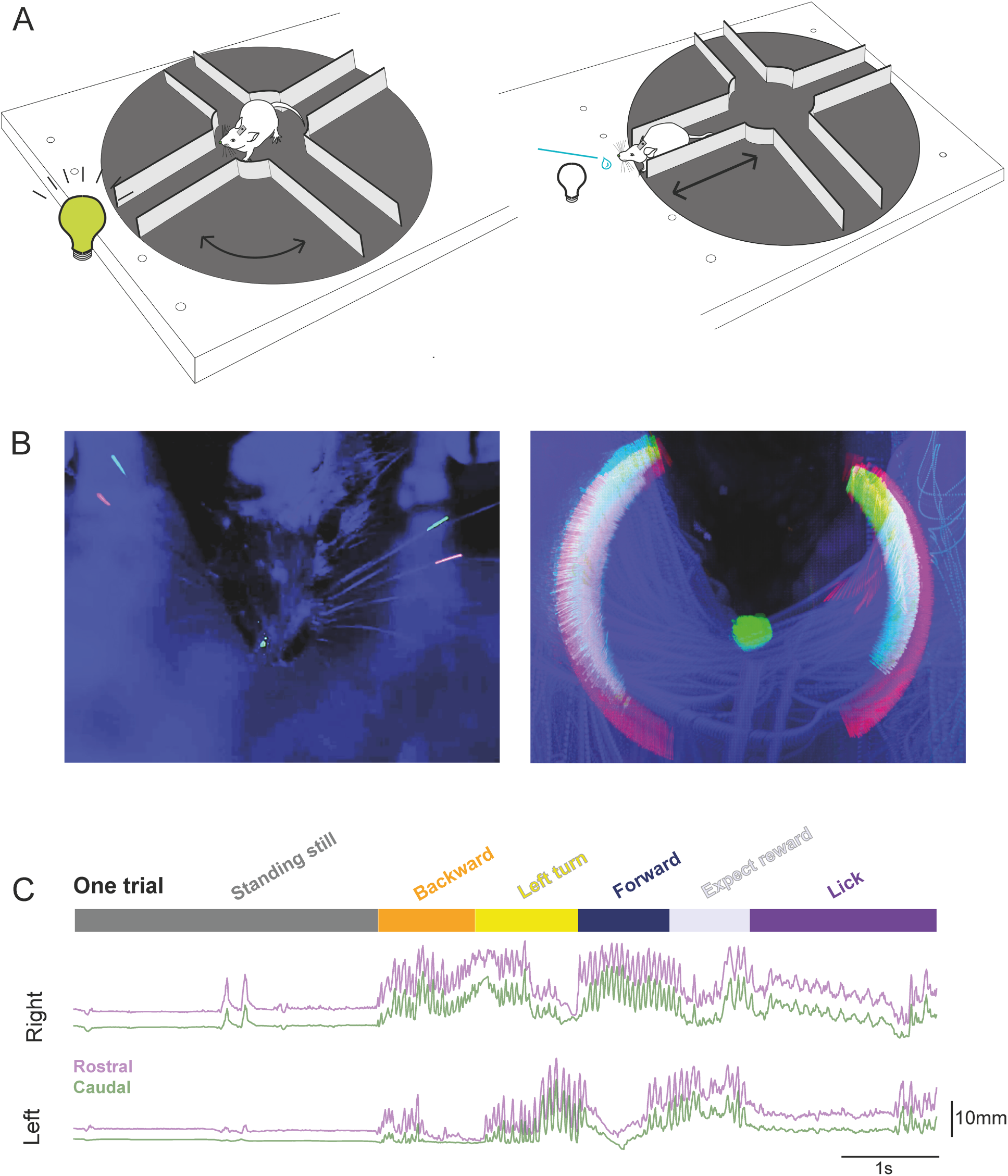
A trial in a “Air-track” plus maze with whisker tracking. **A.** *Mice in a plus maze.* A schematic of the air track plus maze used for this study is shown here with a head fixed mouse, navigating the maze. Mice were trained to rotate the maze away from a lane that had a LED light and kept rotating the maze around themselves to find the dark lane, which they entered, and traversed to the end, in order to obtain a milk reward. Note that the mouse, the LED, and the location of the reward remained fixed in place, all that moved was the maze. **B.** *Maximum intensity projection of a mouse, its painted whiskers and nose*. **Left.** The painted whiskers on each side of the face, and a spot painted on the nose are shown here in a single frame. Some of the adjacent, un-trimmed whiskers are clearly visible here. **Right.** Maximum intensity projection of 30 trials. The movement of the whiskers around the face of the mouse forms a halo composed of the two colors painted on the whiskers and reflects all the positions the two painted whiskers occupied in the course of 30 trials in the maze. Furthermore, the small spot painted on the nose transforms into a large spot over the course of the thirty trials, indicating that the nose moves a lot as the animal traverses the maze. **C.** *A single decomposed trial of behavior and bilateral whisker motion*. We tracked the motion of two whiskers (rostral one painted red, caudal one painted green) bilaterally for the duration of each trial. Once a trial ended, the lick tube was retracted, LED light turned on indicating that the mouse was in the wrong lane. At this point, mice often waited at the end of the lane without moving much (standing still– dark grey bar on top) for seconds to minutes, before they exited the lane by going backward (orange). Then the mouse turned and rotated the maze around itself (yellow), until the LED at the end turned off. Mice had to enter the dark lane, move forward into it (dark blue). At the end of the lane they waited for lick tube to descend (expect reward – light grey), and lick for the reward (purple). There was a delay between the end of the forward motion, and the time for the lick tube to descend to the animal – reward expectation. Two whiskers on each side of the face – one painted red (rostral whisker), and the other painted green (caudal whisker) -- could be tracked for the extent of the trial. The position of the whiskers over the course of the trial was related to the behavioral epochs. Note that whisker motion was related to the animal’s motion. Whisking was apparent when the animal was moving backward or rotating the maze or going forward. In contrast when the animal was standing still whiskers did not move much. And when the animal was licking the reward, the pattern of whisker motion was distinct – low amplitude, rhythmic, bilaterally symmetric -- compared to epochs where the mouse was moving the maze.

A pixy camera / Arduino interface tracked the Air-floating maze position with 35 fps resolution. This interface was used to trigger an actuator that moved the reward spout toward the animal (**Video 1**), when the animal entered the correct lane. This interface was also used to trigger reward delivery.

### Training

A week after surgical preparation, animals were gradually habituated to handling and head fixation on the plus maze. Mice were kept under water restriction and were monitored daily to ensure a stable body weight not less than 85% of their initial starting weight. Habituation consisted of head fixing mice in the plus maze, while manually giving condensed milk as a reward. Subsequently, over the next few days animals were guided manually with experimenter nudging and moving the maze under the mouse. Mice learned to rotate the maze, go forward and backward, collecting a reward at the end of a specific lane. The rewarded lane was indicated by an LED (placed at the end of the lane) turning off once the animals were facing the correct lane (Figure 1A).

A single complete trial started when the LED turned on, the animal walked backwards to the center of the maze, rotated the maze, orienting itself to the correct lane (indicated by the LED turning off). Then the animal moved forward to the end of this lane, waited for the lick tube to descend and licked the tube for a reward (**Video 1,2**). Note that a trial could last indefinitely, there was no requirement for the animal to move the maze quickly, or even to keep the maze moving. Thus, individual trials could vary widely in their duration.

Data acquisition began once animals performed ~50 trials in a day. Each day, the same two whiskers on each side of the face and a small spot on the nose were painted red or green using UV Glow 95 body paint (F**igure 1B, left**). The tips of the tracked whiskers were trimmed to ensure that they stayed within the 4 cm width of the lane. All other whiskers were left intact. High speed video was acquired at 190 Hz, with a Basler Camera while the set up was illuminated with two dark lamps. Data was acquired in dark light conditions, where the UV glow colors on whiskers were most clearly visible and distinguishable from the background (Figure 1B).

### Data selection and image analysis

We used 3 days of data from 5 well trained animals. These mice performed 50-100 trials in a single hour-long session. Data used here was from animals that could move the maze smoothly. Trials were selected for analysis if the whiskers were visible, the paint was glowing uniformly, and if in the course of the trial, the view of the whiskers was not obstructed by the motion of the animal (Nashaat et al., 2017).

Behavioral states were annotated manually by marking the frames when state transitions occurred. The time point of entry into or exit from the lane was determined by using the position of the eyes, in relationship to the edges of the lanes. The frame on which the animal started moving continuously in one direction was defined as the onset of forward or backward movement.

Data were acquired as Matrox-format video files, each file covering a single trial. These files were converted into H.264-format using ZR view (custom software made by Robert Zollner, Eichenau, Germany). Maximum intensity projections of painted whiskers and nose position within one session were created using Image J. From the maximum intensity projection, three individual rectangular regions of interest (ROI) were selected using the rectangular selection tool in ImageJ. Two ROIs included whiskers, on each side of the face, and a third one was set around the nose (Figure 1B, **right**). The ROI dimensions were calculated using the Measurement tools in ImageJ.

### Tracking

Whisker and nose position were tracked for each session, and for each animal separately. A custom-made ImageJ plugin (https://github.com/gwappa/Pixylator, version 0.5) or an equivalent Python code (https://github.com/gwappa/python-videobatch, version 1.0) was used to track the pixels inside the ROI selected with ImageJ. For each frame, the pixels that belonged to a particular hue value (red or green) were collected and the center of mass for the pixels was computed using the brightness / intensity of the pixel. If tracking failed in some frames, i.e. the algorithm failed to detect any matched pixels – because of shadows, movement, or the whisker getting bent under the animal or against a wall – these frames were dropped and linear interpolation was used to ascribe position values in the missing frames. From this analysis, we created masks for each whisker (**Video 3**), that tracked the whisker position for the entire session.

### Analysis

The following analytical procedures were performed using Python (https://www.python.org/, version 3.7.2), along with several standard modules for scientific data analysis (NumPy, https://www.numpy.org/, version 1.15.4; Scipy, https://www.scipy.org/, version 1.2.0; matplotlib, https://matplotlib.org/, 3.0.2; pandas, https://pandas.pydata.org/, 0.23.4; scikit-learn, https://scikit-learn.org/stable/, version 0.20.2). Asymmetry of whisker position was assessed on a trial by trial basis, using values of normalized positions, which made it possible to compare side to side differences in position. In each trial, the most retracted and the most protracted positions became 0 and 1, respectively. The “∆R-L” values were computed by subtracting the left whisker value from its right counterpart on each time point. Thus the ∆R-L value ranged from −1 to 1: a value of “1” meant that the right whisker protracted maximally while the left whisker retracted maximally (i.e. the whiskers orient fully leftward). A value of “0” indicated that the left and the right whiskers protract/retract to the same extent. In order to test asymmetry, a Wilcoxon’s signed-rank test was used to test for the left vs. right normalized positions.

The nose position was normalized in the same manner as whisker position, with the exception that for the nose, the right-most and left-most positions become −1 and 1, respectively.

Normalization of duration for each behavioral state was performed by resampling. For each epoch, we fitted the (normalized) time base from −0.2 to 1.2 with steps being 0.01, where the epoch starts at time 0 and ends at time 1. The data points were resampled from the original time base (i.e. frames) to the normalized time base, using interpolation. The normalized data could then be used for calculating averages and standard error of mean (SEM) of whisker position in the course of an epoch. We used a Wilcoxon signed rank test to assess whether whiskers were asymmetrically positioned in different behavioral states. We computed the R^2^ values and used the Kruskal–Wallis test to assess whether the position of whiskers on each side of the face, or the asymmetry of whisker position best captured the variance in side to side movement of the nose.

### Calculation of whisking parameters

Three whisking parameters were calculated from the whisker position traces: set point, amplitude of whisking, and frequency. Set points of whisker position and amplitude of whisking were computed directly from whisker position traces, using a 200 ms sliding window around each time point. Within this window, the “set point” was defined as the minimum (the most retracted) value while the “amplitude” was defined as the difference between the maximum (the most protracted) and the minimum values (the set-point). The sliding-window algorithm was based on the Bottleneck python module (https://github.com/kwgoodman/bottleneck, version 1.2.1).

The frequency components were estimated from time-varying power spectra obtained through wavelet transformation (Morlet wavelet where the frequency constant was set at 6 for which we used the “wavelets” python module: https://github.com/aaren/wavelets, commit a213d7c3). The power spectra data between 5-35 Hz derived from each animal was pooled (movement data from both sides was used). Three frequency components were estimated using a non-negative matrix factorization (NMF). Here the “decomposition.NMF” class of the scikit-learn module, without L1 regularization was used. The modal-values of the low, the medium and the high-frequency components for each animal were ~ 5–8, 8–15 and 15–25 Hz, respectively. The power of each frequency component was defined as the coefficient of the component of the power spectrum at a given time point, multiplied by the power of the component.

### Classification analysis

For classification and decoding, we used 5 motion parameters taken from both the left and the right whiskers (10 parameters in total): the whisking set point, amplitude, and the power of the 3 frequency components. Before inputting these parameters to the classifier, the right and left whisker parameters were mixed and transformed into “offset” and “asymmetry” parameters. The offset and asymmetry parameters were computed for each of the five whisking parameters (referred to in general as variable X here). The values for each side X_Left_ and X_Right_ were computed for each time point, and these values in turn were used to derive the offset value X_offset_ and an asymmetry value X_Asymmetry_, using the following equations (in case of the left-turning animals): X_offset_ = (X_Left_ + X_Right_)/2, and X_Asymmetry_ = (X_Left_ - X_Right_)/2. For right-turning animals, the sign of X_Asymmetry_ was inverted.

We used naïve-Bayes classifiers with Gaussian priors that predicted the animal’s behavioral state based on the baseline and the asymmetry values of the five parameters of whisker motion. The “naïve Bayes.GaussianNB” class of scikit-learn was used with default parameter settings. Because the labeling of whiskers varied across sessions, a classifier was trained to predict the behavior of each animal during each daily session. For training the classifier, the ten parameters were first pooled for all trials during a session. To avoid selecting periods where the animal was transitioning from one epoch to another, we pooled time points from the midpoint of an epoch and including just the middle 80% of each behavioral epoch. Classifiers were trained with 500 randomly chosen timepoints for each epoch. In most cases, single timepoints were not resampled during the selection process; the only exception was if the total number of timepoints for the epoch was less than 500. This was an issue in two sessions, one for a *Forward* epoch, and another for a *Expect-reward* epoch.

The accuracy of the classifier was tested by comparing predictions against manual annotation. First, we selected parameter sets from another 200 randomly chosen timepoints from each behavioral epoch; these epochs could overlap with those used for training the classifier. Then the prediction of the classifier was tested against the manual annotation. Finally, the probability for each behavioral state was computed using the “predict_proba” method of the “GaussianNB” class. We used twenty iterations of training and testing a classifier for each session i.e. randomly picking up 500 and 200 timepoints per epoch for testing and training, respectively. The scores of the 20 classifiers built to classify one session were averaged.

One caveat in performing this classification was that the whisker motion changed dramatically in the course of each behavioral epoch; it was different in the first and second halves of many behavioral epochs. To avoid underfitting, the classifier was trained to predict “half-epochs” (e.g. Standing-still 1st half, Standing-still 2nd half, Backward 1st half, etc). The output from the classifier was then merged into full-epochs: a classifier that returned “Turn 1st half” as an output was said to have categorized the input parameter set to be in the Turn state. Note that the probability of being in the Turn state was computed by summing the probabilities, for the “Turn 1st half” and the “Turn 2nd half” half-epochs.

The contribution of each parameter to accuracy of classifiers was calculated by randomizing a subset of parameters. To generate a completely shuffled dataset the manual annotations of behavioral epochs in the training dataset were shuffled. The contributions of baseline and asymmetry values were estimated by separately shuffling all 5 baseline or 5 asymmetry values -- including set points, amplitudes, and 3 frequency components -- in the training dataset. Contributions of the set point and the amplitude were separately estimated by shuffling the baseline and asymmetry values of the corresponding parameters. For estimating the contribution of frequency, the values of the three frequency components (baseline and asymmetry) were shuffled separately and the resultant reductions in accuracy were considered as contributions to accuracy of the classifiers.

## Results

### A single trial in the Air-track plus maze

The behavior in the maze was simple, self-initiated, and had no time constraints. Mice decided when to begin a trial, and how long to spend on a trial (Figure 1C, **Videos 1, 2**). At the beginning of a trial, when mice were at the end of a lane, just after they had obtained a reward, the LED was turned on. Mice then had to move backward out of the lane to reach the center of the maze. When they reached the center of the maze, they turned it around themselves to find a dark lane, a lane in which the LED light was off. Mice then entered this lane – moved forward in it, all the way to end, where they waited for the reward tube to descend down toward them and then they licked the tube for the milk reward. When mice entered incorrect lanes, i.e. the lane where the LED light was on, they were not rewarded and had to move out of the incorrect lane, rotate the maze, and find the dark lane (**Videos 1, 2**). Over the course of training, mice learned to steer the maze and to select the correct lane quickly in each trial. During navigation, the behavior of mice could be divided into distinct states, consistent enough to be classified into epochs (Figure 1C): (1) the “Standing still” epoch that marked the beginning of a new trial, where the mouse stood still at the end of a lane, typically after previous reward delivery; (2) the “Backward” epoch, where the animal moved backward out of the lane; (3) the “Turn” epoch, where the animal entered the center of the maze, and rotated the maze left or right around itself until it chose a new lane; (4) the “Forward” epoch, where the mouse moved forward into the new lane; (5) the “Expect reward” epoch, when the mouse waited at the end of a lane for the reward tube to descend; (6) the “Lick” epoch, if the chosen lane was correct, the reward tube descended towards the animal. This epoch lasted until the tube was retracted up and away from the animal.

### Stereotypical whisking in each behavioral epoch

We tracked the movement of C1 and Gamma whiskers as mice (n = 5 animals) navigated the plus maze (**Video 1-3**). Whiskers on one side of the face in general moved to the same extent, and they were at similar set points with respect to each other in all behavioral epochs over the course of a trial. Therefore, for the rest of our analysis, we compare only the motion of a single whisker the rostral C1 whisker that was painted red, on each side of the face (Figure 1B).

In general whisker motion showed four stereotypical characteristics in different behavioral stages (**Video 4**): 1) When mice moved in the maze, large amplitude (high frequency) whisking with a high degree of asymmetry between the sides of the face was evident (i.e. in backward, forward, or turning epochs) (Figure 1C, **2**). 2) In contrast, when mice were simply standing still in the maze, whisker movement was negligible and the set point of whiskers was retracted in this epoch (Standing still, Figure 1C, 2). 3) When mice were standing still but were licking the reward tube, whisking was regular, and occurred at lower frequency than when mice were moving. When mice were standing still, but were licking, the set-point for whiskers was also protracted compared to times when mice were at the end of lane and standing still (Figure 2). 4) Finally, during reward expectation, whiskers were protracted and showed high amplitude, and high-frequency whisking.

**Figure 2.**
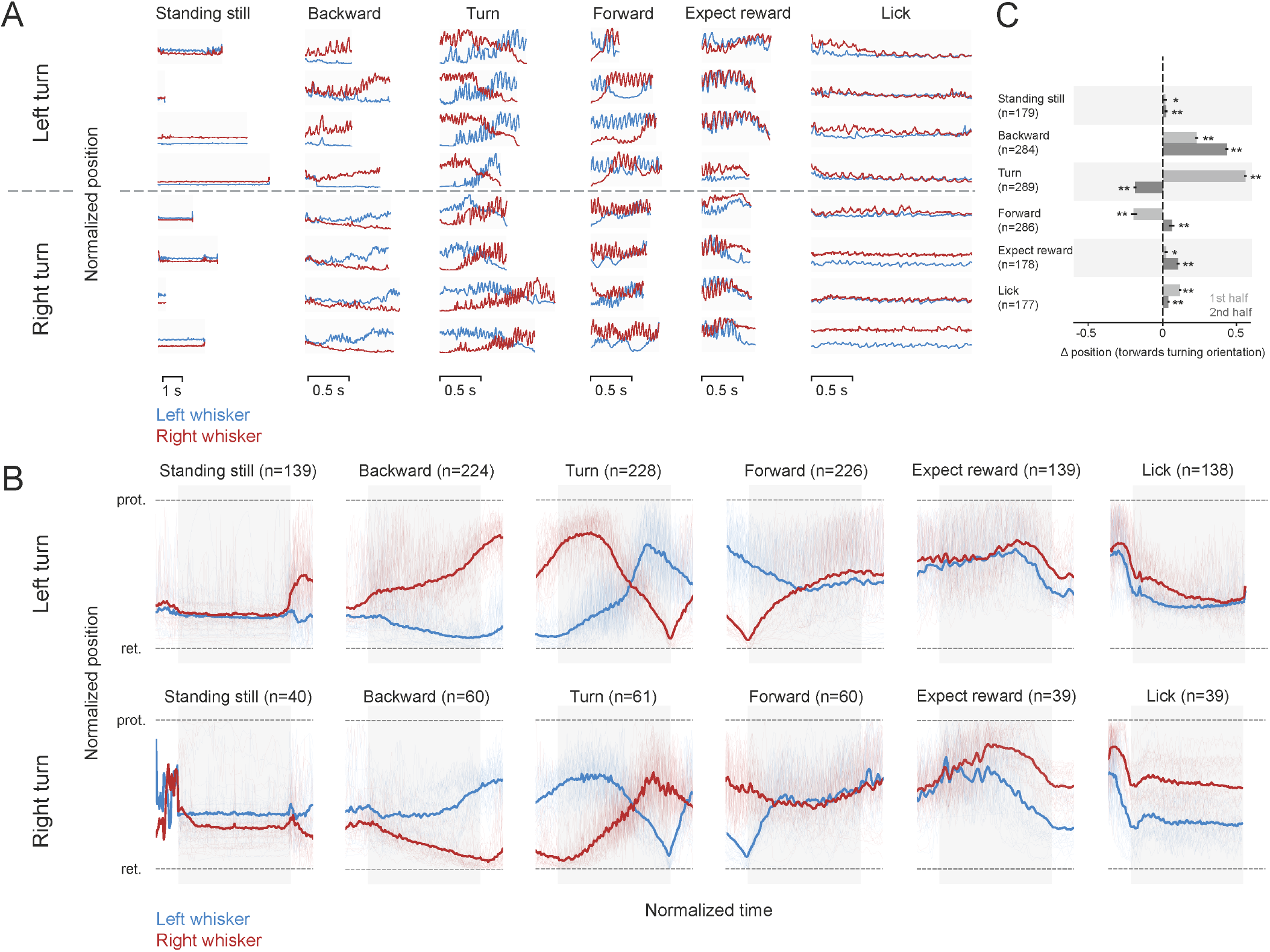
Patterns of whisker motion during different behavioral epochs. **A.** *Normalized whisker position in single trials and behavioral epochs*. The position of whiskers on each side of the face (red is right side, blue is the left side) in the course of each behavioral epoch (without averaging or smoothing), taken on different days are shown from 3 ***left*** turning animals (top, above the dashed line), and 2 ***right*** turning animals (below the hashed line). A trial was decomposed into behavioral epochs selected for analysis. **B**. *Normalized average whisker position in different behavioral epochs.* Whisker asymmetry in different behavioral epochs averaged during behavior for left turning (top), and right turning (bottom) mice. Normalized whisker position on left and right side, for left-turning (n=3) and right-turning (n=2) animals. While whiskers were retracted almost equally on both sides of the face, before the trial began the average normalized data reveal that as mice begin to go backward whisker positioning becomes asymmetric and the asymmetry was related to the direction of turn mice impose on the maze. When mice moved the maze to the right (bottom panels), the positioning of whiskers was asymmetric, and a mirror image of how whiskers were positioned when mice turned the platform to the left (top panels). The blurred red and blue traces in the background show whisker positions in every single trial. The epochs used for analysis are defined by the grey bars behind the average whisker position traces. **C.** *Average whisker position in first and second half of behavioral epoch*. The average side to side position of whiskers changed significantly in all behavioral epochs (Wilcoxon sign rank test, p < 0.05 single asterisk and p < 0.01 double asterisk, combined for right and left turning trials). The normalized mean asymmetry for the first half of each epoch was 0.02 (SEM 0.01) for standing still, 0.23 (SEM 0.01) for backward, 0.56 (SEM 0.01) for turn, −0.2 (SEM 0.02) for forward, 0.023 (SEM 0.01) for expect reward, 0.12 (SEM 0.01) for lick. The normalized mean asymmetry for the second half of each epoch was 0.02 (SEM 0.01) for standing still, 0.432 (SEM0.01) for backward, for −0.19 (SEM 0.01) turn, 0.06 (SEM 0.02) for forward, 0.1 (SEM 0.01) for expect reward, and 0.04 (SEM 0.06) for lick. In the right turning animals (data is not shown), the mean values were −0.65 (SEM 0.02) for standing still, −0.14 (SEM 0.02) backward, −0.46 (SEM 0.01) for turn, 0.08 (SEM 0.02) forward, 0.08 (0.02) expect reward, and 0.27 (0.03) lick in the first half and −0.09 (SEM 0.02) stand still, −0.31 (0.02) backward, 0.119 (0.02) turn, −0.03 (0.03) forward, 0.28 (0.03) expect reward, and 0.26 (0.04) lick. The light red and blue traces show the single trial data that was used to compute the average whisker positions on two sides of the face. The epochs used for analysis are defined by the grey bar.

Whisking was so stereotyped, that it served as a “signature” of each behavioral epoch (Figure 2). Even though the duration of a behavior, i.e. standing still or moving backward varied from trial to trial, the positioning of whiskers bilaterally reflected the behavior.

The whisker position traces from each trial were averaged after normalizing for time and bilateral extent of whisker movement (in order to compare whisker position on the two sides of the face) (Figure 2B). Averaging smoothed out the rhythmic whisking motion i.e. averaging removed the fast components of whisking. These averages confirmed what was evident in the raw data. Each behavioral epoch thus had its own whisking signature and this was independent of animals, sessions and trials.

### Bilateral asymmetry signals turn direction

Mice had a strong rotation preference; they rotated the maze left or right, and rarely turned the maze ambidextrously in both directions. The mouse’s decision to turn right or left was reflected in the asymmetric positioning of whiskers throughout a trial, which was evident very early during the backward movement epochs (n= 284; 224 left turning epochs + 60 right turning). The asymmetry increased as the animal reached the center of the maze. In all trials, when mice reached the center of the maze, animals that turned left retracted their whiskers on the left side to their full extent while simultaneously protracting whiskers on the right side (Figure 2A, B top panels & **Video 1-2**). In contrast, animals that turned right retracted their right whiskers, and protracted their left ones, as they reached the center of the maze (Figure 2A bottom panels). Whiskers on the side of the face that were protracted, showed high amplitude whisking motion, while whiskers on the other side of the face retracted but displayed almost no rhythmic whisking motion (Figure 2 and **Video 1-2**). The asymmetry that began at or just before the onset of backward movement, increased while the animal was approaching the center of the maze and flipped during the turn (Figure 2). There was an inversion in the whisker position as the animals moved forward into a lane: whiskers that were completely protracted during backward movement were retracted, and vice versa. This inversion in whisker position was stereotypic, occurring automatically in every turn epoch (n= 289; 228 left turning epochs + 61 right turning). Note that mice maintained this asymmetric position as they moved forward into the lane, primarily as a result of whisker contact with the wall on one side of their face. The asymmetry gradually diminished as mice moved further forward into the lane (n= 286 forward; 226 left + 60 right).

To quantify these changes in asymmetry in the course of each behavioral epoch, we divided each behavioral epoch in two and compared the normalized left whisker position to the right whisker position (∆R–L) at the beginning and end of each behavioral epoch. In all behavioral epochs, in both right and left turning animals, there was significant whisker asymmetry (p < 0.05, Wilcoxon sign rank test, n =139 to 228 epochs for left turning animals, and n = 40 to 61 epochs for right turning animals, Figure 2C) and asymmetric positioning changed significantly (p < 0.01, Wilcoxon sign rank test) when whisker asymmetry in the first and second half of each behavioral epoch were compared (Figure 2C, data for right and left turning animals are binned together). There is a caveat to note here: the small difference in side to side positioning of whiskers during reward expectation, and licking arose in part from the direction of descent of the lick tube (**Videos 1, 2**). Independently of whether mice propelled the maze right or left, the reward tube descended toward the left side of their face, consequently, during licking and reward expectation the side to side asymmetry in all animals shows a similar pattern (compare the whisker position in reward expectation and licking epochs for the left and right turning mice in Figure 2B during reward expectation and lick epochs). By the time the animal finished licking the reward tube, and decided to move backward in the lane, the lick tube related asymmetry was no longer evident; instead a small but consistent and significant (p < 0.05, Wilcoxon sign rank test) difference in side to side to position of whiskers was evident (compare the left and right turning data, in standing still epochs in Figure 2A, B). Taken together our results suggest that whisker asymmetry is a constant feature, and whiskers are actively repositioned as mice move through the maze. Whisker position at the beginning of a trial can predict decisions mice make in imposing a movement direction on the maze, and the extent and direction of the asymmetry can effectively map the position of the animal in the maze.

In principle, whisker position asymmetry could arise from changes in amplitude or frequency of whisking or changes in set point (Figure 3). To examine these possibilities, the time varying power spectra of whisking, and the amplitude of whisking and the set point of whisker position were calculated. The time varying power spectra revealed multiple frequency bands, one band that spanned 0-8 Hz, another that spanned 12-30 Hz and an intermediate band from 8-12 Hz (Figure 3A-3D). When mice were standing still they did not whisk much, as they began moving backward whisking frequency was low (Figure 3C), but as they continued moving the frequency components in whisking increased.

**Figure 3.**
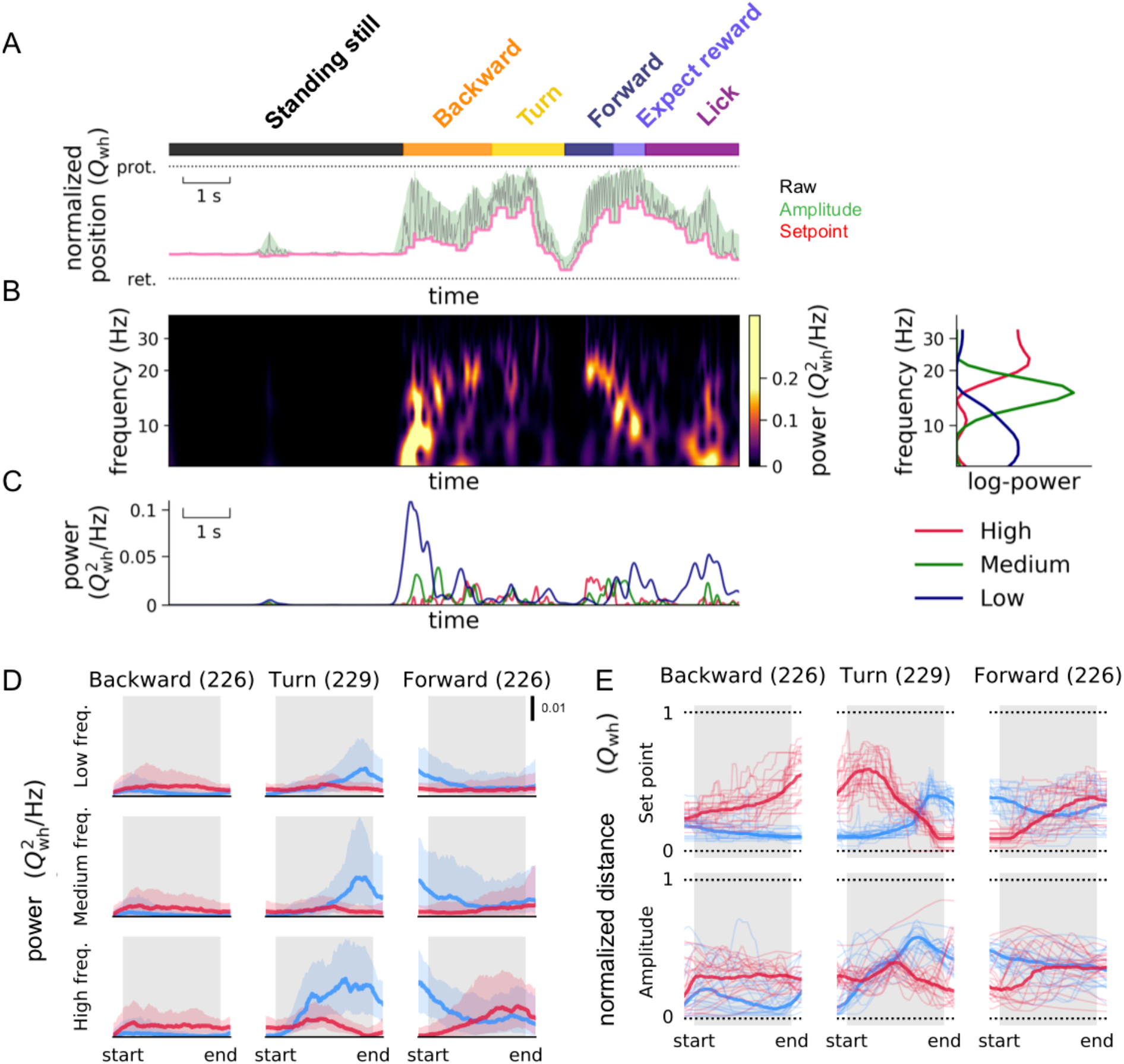
Behavioral state dependency and whisking frequency, set point and amplitude. **A**) *Extraction of set point and amplitude of whisking.* The behavioral epochs -- standing still (black rectangle), backward (yellow rectangle) etc -- were annotated in the same way as in Figure 1. A 200 ms-sliding window was applied to the whisking trace (grey) to detect the set points (minima in the window; pink line) and amplitudes (difference between maxima and minima; green) of whisking. The normalized whisker positions (*Q*_*wh*_) were plotted with the most retracted (ret) position set to zero, and the most protracted (prot) whisker position set to one. (**B**) *Extraction of time-varying power spectra of whisker motion*. The time varying power spectra of whisker motion were calculated (left), and 3 frequency bands evident in the power spectra -- high-frequency (the modes typically locaed at 12-30 Hz, red), medium-frequency (8-12 Hz modes, green) and the low-frequency (0-8 Hz modes, blue) components --were plotted on the right. (**C**) *Behavioral state and time varying frequency components*. The 3 time varying frequency components plotted over the course of a trial show that for a single trial plotted here, the high frequency components were more common as mice finished turning, and the low frequencies were common as the mouse began to move backward. (**D**) *Set point of whiskers, and amplitude of whisker motion during different behavioral states*. When mice were moving backward, and or turning, asymmetry in set points and amplitude developed differently over time (compare middle and bottom plots, the whisker position plots in the top row are plotted here as reference, taken from Figure 2). Thick lines are median values. Thin lines in show the values from 20 representative epochs across all sessions from different animals (n=5 sessions from 3 left-turning animals; numbers in brackets refer to total number of epochs analyzed). In these behavioral epochs, the asymmetry in whisker positions can be attributed to asymmetry in set points. The normalized mean asymmetry for *set point* of whisker position in the first half of each epoch was 0.17 (SEM 0.006) for backward, 0.47 (SEM 0.007) for turn, −0.13 (SEM 0.01) for forward motion. The asymmetry for the set point in the second half of each epoch was 0.31 (SEM 0.007), −0.03 (SEM 0.006), 0.05 (SEM 0.01) for backward, turn, and forward motion respectively. Similarly, asymmetry for the *amplitude* of whisking in the first half of each epoch was 0.12 (SEM 0.008), 0.01 (SEM 0.007), −0.12 (SEM 0.02) for backward, turn, and forward motion. Asymmetry in *amplitude* of whisking for second half of each epoch was 0.22 (SEM 0.007), for −0.24 (SEM 0.006), and −0.035 (SEM 0.01) for backward, turn, and forward motion. (**E**). *Whisking frequency on right and left side during different behavioral states*. The side to side difference in power in the 3 frequency components of whisker-motion was highest as animals turned the maze, and as they entered a new lane. The shaded regions show the 25 and 75 percentiles of the values across all sessions from different animals (n=5 sessions from 3 left-turning animals; numbers in brackets refer to total number of epochs analyzed). The epochs used for analysis are defined by the grey bars in the background. Data shown here is from left-turning animals.

The amplitude of whisking (Figure 3A, **green**) and the set point (Figure 3A, **pink**) both reflected asymmetry. In the course of the backward movement (n= 224 left turning epochs) and the turn (n=228), the set point and amplitude traces showed distinct patterns suggesting that these aspects of whisking change independently during movement in the maze (Figure 3E). When mice moved backward, the asymmetry in set point continued to increase, i.e. mice protracted one side more. When mice turn, the set point and amplitude of whisking changed at different times in the behavior (Figure 3E). When mice move forward (n= 226 left turning epochs) into a lane both the set point and amplitude of whisking change at the same time and in the same direction (Figure 3E, **Right panel**). Taken together these results suggest that the bilateral amplitude of whisking and the set point of whisker position are actively and independently controlled, and independently contribute to asymmetry.

### Decoding behavioral state from whisking parameters

The behaviors in the plus maze are complex: mice can move rapidly or slowly; behavioral epochs can be interrupted by pauses when the mouse takes a time out from the behavior. The time taken to complete a trial can vary widely. Nevertheless, because whisking in each epoch is so stereotyped – every trial shows the same dynamics of whisker movement – it is possible to use Bayesian analysis to decode the behavior of the mouse from the whisking parameters, i.e. frequency, amplitude and set point. To examine whether behavioral state could be predicted from one or more whisking parameters we used a naïve Bayes classifier, where each feature of whisking -- set point, amplitude and frequency -- was examined independently. Each parameters contribution to the correct classification of the behavioral state was assessed (Figure 4A). For a single trial, the probability of correctly classifying each behavioral state from whisking parameters was high (Figure 4B). But note behaviors overlap with each other: when the mouse is “*Licking*” it is also “*Standing still*”, when the mouse is *“Turning”* it is also going “*Forward”* or *“Backward”*. When behaviors overlap, whisking parameters also overlap and make it a little bit harder to correctly classify behavior from the whisking parameters. When we trained the classifier and used it on the dataset (left matrix), and on the shuffled data set, behavioral states could be accurately classified 80% of the time (Figure 4C), with a 20 % correct classification even after shuffling the data – i.e. there was a baseline effect of 20% (Figure 4D). *These data suggest that whisking parameters can predict where the mouse is, and what the mouse is doing in a maze*.

**Figure 4.**
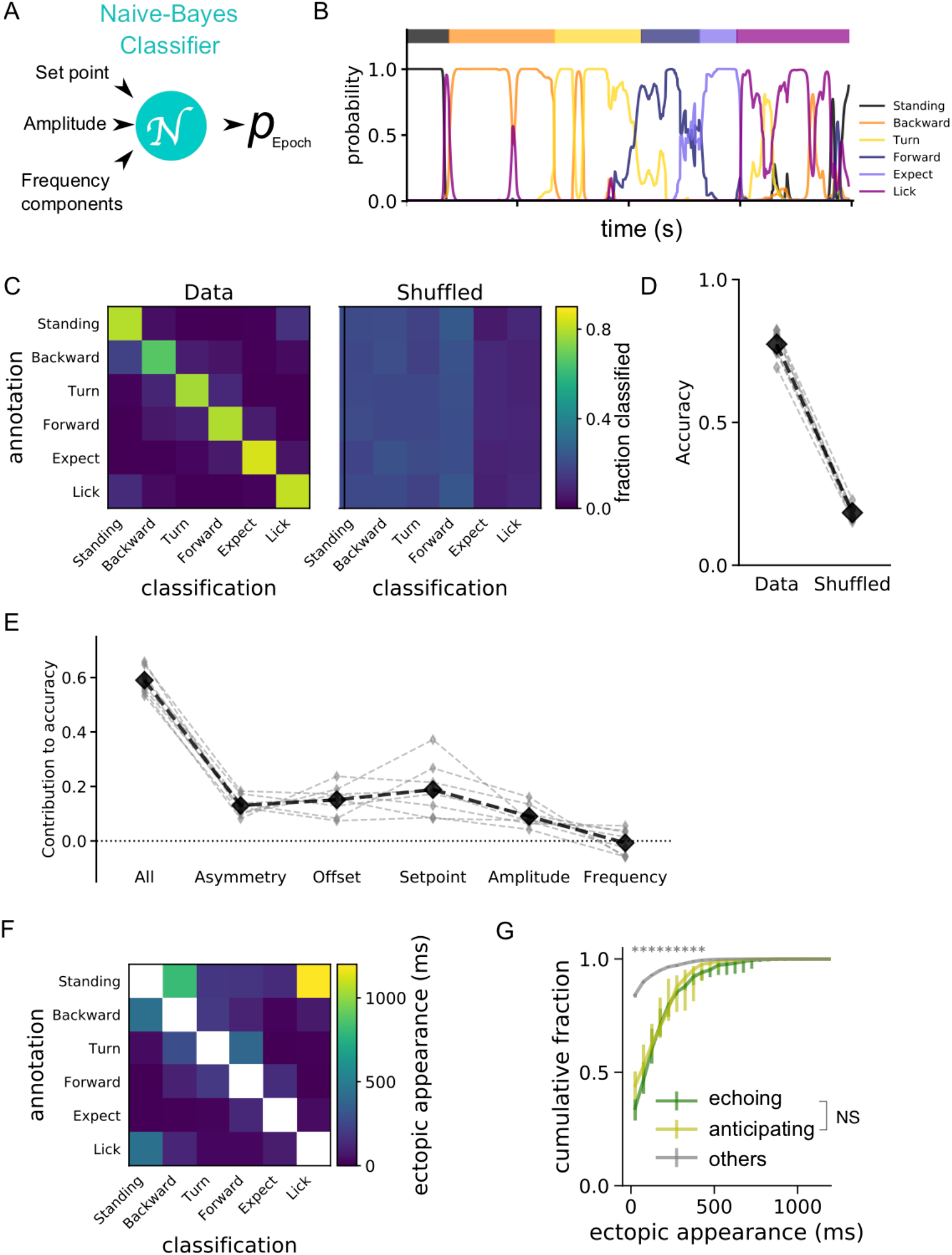
Classification, decoding and prediction of relationship between behavior state and whisking. (A) A naïve-Bayes classifier with Gaussian priors was used to output the probability of a specific behavioral state (*p*_*Epoch*_) based on Set point, Amplitude and frequency parameters of whisker motion. In total, 20 distinct Classifiers were trained for each session. (**B**) T*he output of probabilities in a representative trial.* The individual colored lines show the probability output generated by the classifier for each behavioral epoch. The ground-truth behavioral epochs were annotated manually and are shown on the top of the graph. (**C**) *Confusion matrices for the classifiers*. Classifiers trained with the actual dataset (left matrix) correctly classified behavioral states, 80% of the time; those trained with shuffled annotations (right matrix) generated chance-level (20%) accuracy. The rows are the categories of the manually annotated behavioral epochs used for testing the classifiers, and the columns are the categorical output from the classifier. The fraction of time points belonging to each manually annotated behavioral state are represented by the colors of each square. (**D**) *Accuracy of the classifiers*. Gray marks indicate the scores for each session (n=10 sessions from 5 animals), whereas black marks show the averages across sessions. *p=0.0143, Wilcoxon signed-rank test. (**E**) *Contribution of individual parameters to accuracy of classifiers.* Asymmetry, set-point and Amplitude contributed significantly to the ability of classifiers to decode the behavioral state of the animal, whereas whisking frequency generated no additional effect. Each parameter set was randomized individually, and reduction in accuracy was computed for each session. The small grey diamonds show the changes in accuracy in individual sessions, the black diamonds are the averages. Kruskal– Wallis test performed for each combination of parameters, revealed significant effects for asymmetry vs frequency, (*p=0.0363), offset vs frequency (**p=0.0012), set point vs frequency (***p<0.0001), and was p>0.1 for all the other combinations, with the omnibus probability being ***p<0.0001. **F**) *Misclassifications.* The duration / time points where the probability was higher than the chance level were plotted. There was a tendency that behaviorally neighboring epochs get misclassified i.e. there was “ectopic appearance” of whisking patterns related to certain other behavioral epochs. (**G**) *Smooth transitions between behavioral epochs.* The duration of above-chance probability for neighboring epochs was computed and plotted as cumulative histograms. The effects of “Echoing” and “Anticipating” -- the effects of preceding and following behavioral epochs, respectively -- were found to be significantly smaller in bins more than 500 ms compared to all the other types of ectopic epoch appearances (*p<0.05, Kruskal-Wallis test; n=10 sessions from 5 animals). No significant difference was found between echoing and anticipating epoch appearances (NS, p>0.05 for all the time bins).

Next, we examined whether asymmetry, set point, amplitude of whisking, or frequency contributed significantly to the ability of classifiers to decode the behavioral state (Figure 4E). Five parameters of bilateral whisking were examined, the amplitude, setpoint, frequency, asymmetry and offset. Asymmetry was defined as the difference in position of whiskers on two the sides of the face, offset was defined as the average position of whiskers on the two sides of face, the set point was the most retracted position of the whiskers, and the amplitude was the difference in the most retracted and most protracted position of the whiskers. Our analysis indicated that set point, amplitude and offset of whisking significantly contributed to the accuracy of decoding behavioral state. Frequency of whisking had a negligible additional effect on decoding behavioral state – it added very little, once asymmetry of whisker position, the amplitude of whisking, and set point had been taken into account (Figure 4E).

Finally, we looked for ectopic appearances – i.e. unusual occurrences – in the classification by comparing the what we observed in manually annotated behavioral state, to the predicted behavior state. Decoding of behavior relies on whisking parameters. But if the behaviors mix with each other -- i.e. animal stands still during licking, or stops licking in licking epoch, or stands still during a forward or backward motion -- then decoding cannot be perfect. When the animal was *Standing Still*, it could also be licking, when it was in the *Licking* state, it could just be *Standing still*, when it was *Turning* it could also be moving *Forward* into a lane (Figure 4F).

Another issue in decoding arises from the incomplete separation between behavioral epochs. Transitions between behaviors can be incomplete, and there can be echoing effect i.e. earlier behavioral epochs affect subsequent ones or an anticipation effect where the epoch after affects the one before (Figure 4G). These effects of transitions between states could be minimized if the duration of the behavioral epoch was long, at least 500 ms (Figure 4G).

### Sensory input and asymmetry

Whiskers are the primary tactile organs of rodents, and in principal whisker asymmetry could arise from tactile input to the whiskers, especially in the Air-Track system where whisker contact with the walls is a constant factor. To examine whether tactile input from whiskers drives the side to side asymmetry, we trimmed all whiskers off bilaterally and painted the remaining whisker stubs (Figure 5A, **right**). Mice were able to perform the task without their whiskers (Figure 5B). Furthermore, in these mice, whisker asymmetry still predicted the direction that the animal would move in, and whisker position still varied in a behavioral state dependent manner. There were still significant differences in direction of asymmetry during backward motion (n= 102 left turning epochs), turning (n= 101 epochs) and forward motion (n= 102 left turning epochs) (p < 0.01 Wilcoxon Sign rank test n = 101, Figure 5C). Note that trimming abolished side to side asymmetry during standing still, licking, and expecting reward. These effects of trimming during licking and reward expectation are in part related to the positioning of the lick tube as it descended towards the left side of the animal. That trimming abolished asymmetry during standing still epoch was probably related to the small initial asymmetry in this epoch. These results rule out the influence of sensory input as the primary cause of stereotypical whisker positioning.

**Figure 5.**
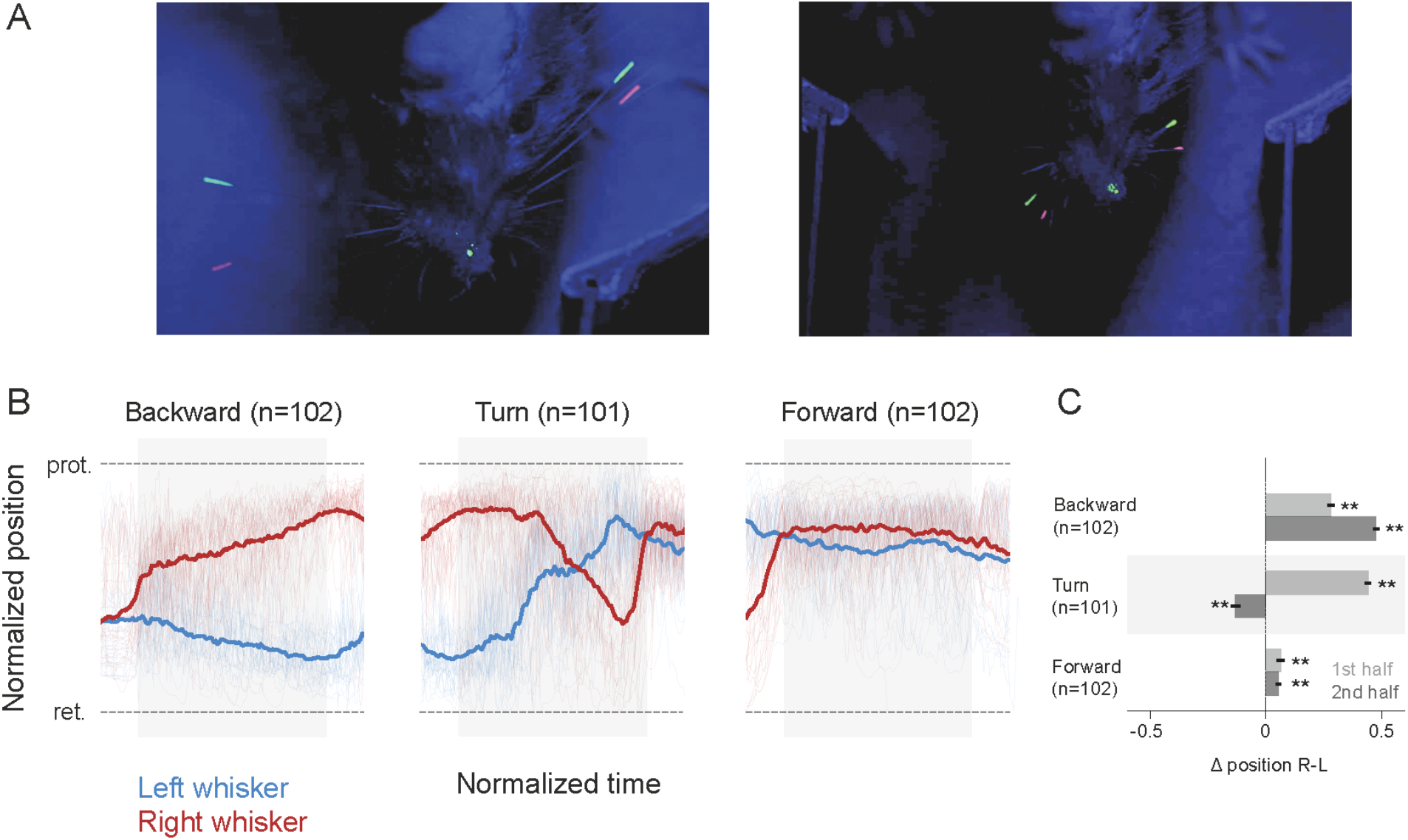
Whisker asymmetry was not related to tactile input. **A.** *Single frames of whisker position in normal (left) and trimmed (right panel).* In left turning mice when whiskers were trimmed, the whisker stubs could still be tracked, side to side asymmetry of whisker position was still evident. **B**. *Normalized average whisker position after trimming in different behavioral epochs.* The asymmetric positioning of whiskers persisted and was evident when mice began to move backward, and persisted through the trial, as mice moved further backward and made left turn. **C**. *Mean asymmetry after trimming*. Even after trimming, there was significant asymmetry (Wilcoxon Sign Rank test, p <0.01, based on n= 102 for backward motion, n=101 turn, n = 102 forward epochs). The normalized mean asymmetry of whisker position in the first half of each epoch was 0.28 (SEM 0.02) for backward, 0.44 (SEM 0.013) for turn, 0.06 (SEM 0.02) for forward motion. The normalized mean asymmetry in the second half of each epoch was 0.48 (SEM 0.01) for backward, for −0.1 (SEM 0.02) turn, 0.06 (SEM 0.01) for forward motion. The light red and blue traces show the single trial data that was used to compute the average whisker positions on two sides of the face. The epochs used for analysis are defined by the grey bars in the background of the whisker position traces.

### Whisker asymmetry in freely moving animals

Earlier work has shown that whisker asymmetry arises and is related to movement of the head (Towal and Hartmann, 2006; Grant et al., 2009; Schroeder and Ritt 2016). These earlier studies were all in freely moving animals. To examine whether the results we obtained in head-fixed mice were an artifact of head fixation, we tracked whisker motion in 2 freely moving animals (Supplementary Figure 1, **Supplementary videos, 5-6**). These animals had been previously trained on the plus maze while they were head-fixed. Whisker tracking in freely moving animals was constrained to just the central portion of the maze, i.e. just as the mouse backed out of a lane, turned and entered another lane. In freely moving mice, head movement made tracking of whisker position difficult. Not only did the head move up and down, mice often rotated their head from side to side. Nevertheless, in a limited set of anecdotal observations, when accounting for head angle, whisker position was asymmetric in the same direction, at the same places, in freely moving animal as in the head fixed mouse (Supplementary Figure 1). Whiskers on the animals left side were retracted, right side protracted as the mouse exited the lane (Supplementary. Figure 1, top panels, backward) and began to turn left (Supplementary Figure 1, **bottom panels, turn**). Once the mouse was in the center of the maze and turning, the whisker position flipped and the mouse protracted whiskers on the right side and retracted them on the left. The white insets in this figure show the position of whiskers for head fixed mice when they were in a similar position in the maze. Head fixed mice showed similar asymmetry to that seen in the freely moving mice.

### Nose movement: relationship to whisker asymmetry

The facial muscles that move whisker pad and whiskers, are controlled by a central pattern generator, that controls breathing and sniffing (Moore et al., 2013; Moore et al., 2014). Some of these muscles also control the motion of the nose (Haidarlu et al., 2012; Haidarlu et al., 2015); in fact the nose and whiskers move in a coordinated fashion (Kurnikova et al., 2017; McElvain et al., 2018). Here we examined whether the nose moved in a behavioral state-dependent manner, and whether nose movement was correlated to whisker motion. As the maximum intensity projection of nose movement over 30 trials shows the nose moved a fair amount in the course of Air-track behaviors (Figure 1B) and moved differently in each behavioral epoch (Figure 6A – bottom trace). Side to side nose movement (i.e. left - right) was best related to the asymmetry of whisker position on the two sides of the face (Figure 6A, middle trace), it was not as nicely related to the motion of whiskers on either side of the face (Figure 6A, top traces, p < 0.001 Kurskal-Wallis test). This becomes especially evident when whisker asymmetry and nose movement were plotted together: bilateral whisker movement (∆position = R-L, purple) and nose movement (green) changed together (Figure 6B). Note that the end of the *Turn* epoch and the beginning of the *Forward* epoch were uniformly associated with whisker contact to the wall which was likely to have interfered with the relation between whisker asymmetry and nose movement. These data show that just as whisker asymmetry is stereotyped in the behavioral epochs, nose movement was also stereotypical. We conclude that the facial expression governed by the contraction of the various muscles in the face all change in a stereotyped fashion in coordination with behavioral sequences mice express in the maze.

**Figure 6.**
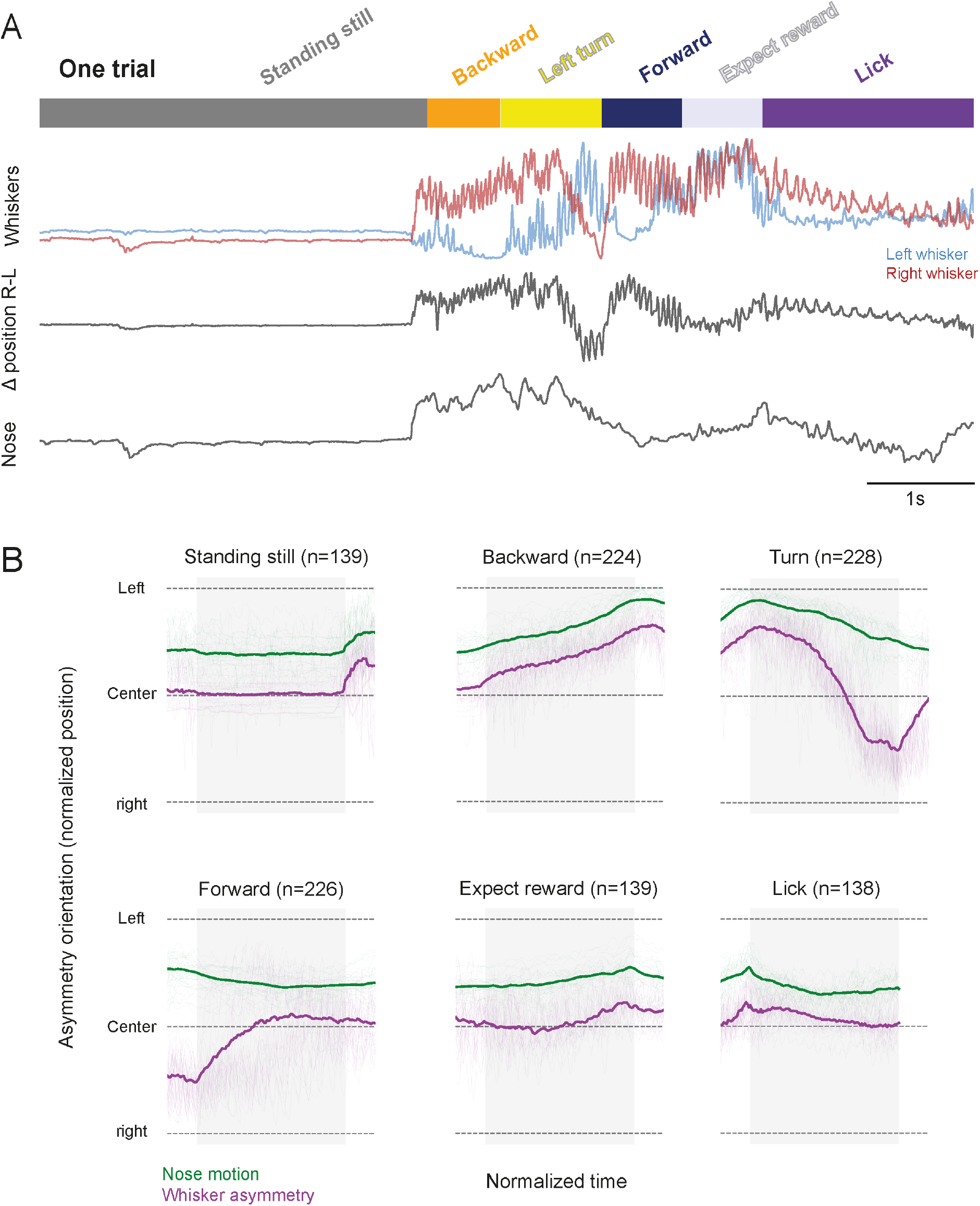
Whisker asymmetry and nose movement. **A**. *The nose moves when mice move*. Whenever the mouse moved its whiskers (top traces red and blue), whisker movement was asymmetric (middle trace) and the mouse moved its nose (third trace from top). The right-left movement of the nose was significantly and best correlated with the complex motion of whiskers (second trace) on both sides of the face (Kurskal Wallis test p <0.001), not with motion of whiskers on any side of the face. **B.** *Average nose movement related to whisker asymmetry in the different behavioral epochs*. When mice are standing still, licking or expecting reward the nose (green trace) does not move and asymmetry in whisker position (purple trace) is minimal but when mice are moving the maze, when mice go backward, turn and go forward the nose also moves and this movement is related to the asymmetric positioning of the whiskers.

## Discussion

Here we show that features of whisking correlate with and can reliably decode the behavioral state of the mouse. Each behavioral epoch has its signature whisking features which made it possible for whisker asymmetry to predict whether the mouse was moving forward or backward, turning, licking, or expecting reward. Even though the Airtrack platform we use here is rich in tactile features like walls and surfaces, and even though whiskers are tactile organs, this relationship between whisking and behavioral state had little to do with tactile input to the mouse. Instead the position of whiskers was related to motor preparation, postural adjustment and active whisking during navigation. Even when mice moved backward in a straight line, the extent and direction of side to side asymmetry in whisker position correlated with the turn that the animal was about to initiate (Figure 2). The signature whisking features that made it possible to relate each behavioral epoch to the positioning of the whiskers (Figure 7) serve as a map of “what the mouse was doing” and “where the mouse was” in the maze. An astonishing aspect of the relationship between whisking and behavior of the mouse is that whisking is not instructed it is not necessary for the task performance, but it occurs spontaneously in the course of the search for the correct lane.

**Figure 7.**
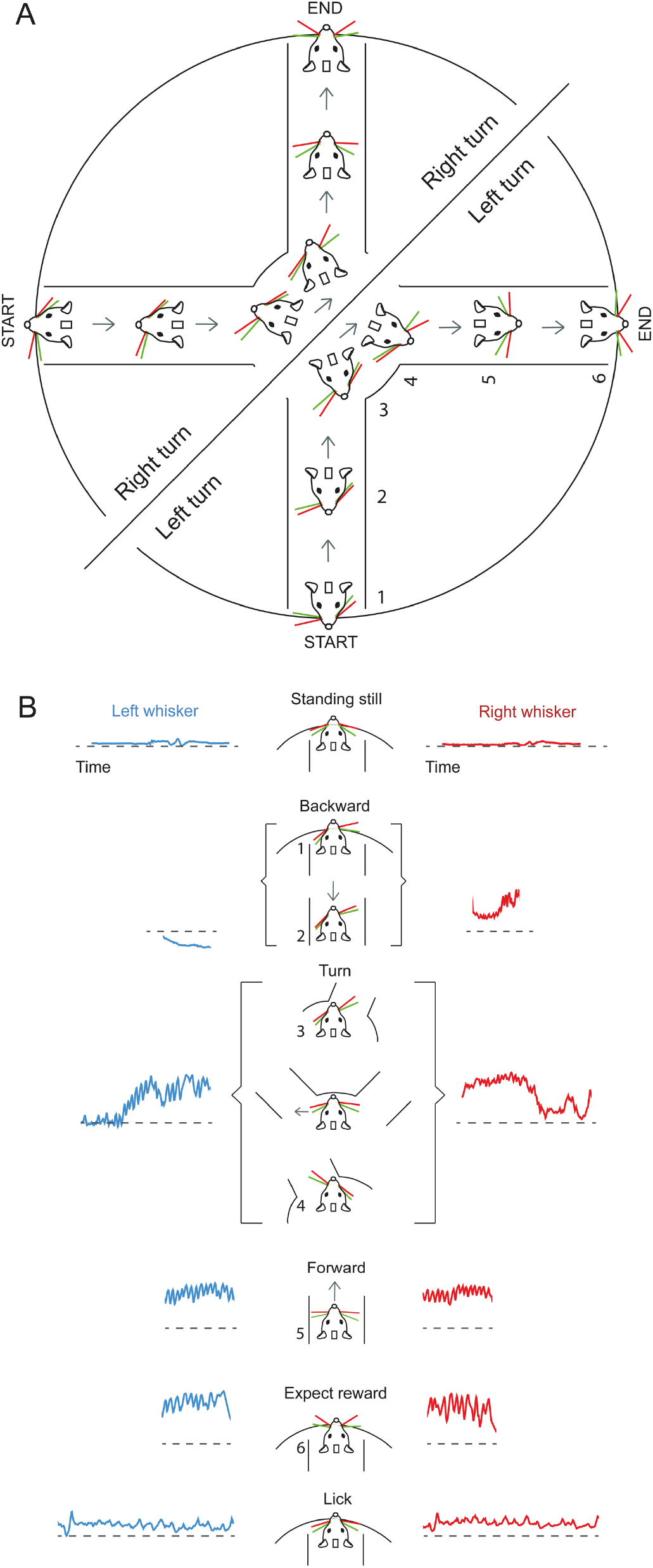
Whisking is a signature of motor plan and behavioral state. **A.** *Right and left turning mice, plan their movement in the plus maze, and this plan is reflected in how they position their whiskers*. Left turning mice protract their whiskers on the right side while moving backward, and right turning mice, protract their whiskers on the left side while moving backward (mice in the semicircles show whisker position for right and left turning mice). The numbers next to the mice indicate the different behavioral epochs. At the onset of backward movement (1) whiskers are already asymmetric. As the animal continues moving backward toward the middle of the maze (2) the asymmetry in whisker position increases until the mouse exits the lane and is in the middle of the maze (3). As the mouse moved forward into a new lane, the asymmetric whisker positioning flipped (4) and then as the mouse proceeded into the lane, the asymmetry diminished (5). The trial ends when the animal is licking (6). **B***. Schematic of whisker position as a signature of behavioral state and movement preparation*. Asymmetry in whisker position and whisker set point both reflect the distinct behavioral state of the animal. The numbers in the schematic are the behavioral states (related to the numbers used in A).

A priori we expected that a portion of the whisker asymmetry would arise from tactile input from the whiskers. After all each whisker is associated with thousands of neurons in the rodent somatosensory cortex (Van der Loos and Woolsey, 1973; Oberlaender et al., 2012), and stimuli to individual whiskers evoke a cortical response (Simons, 1985; Armstrong-James et al., 1992; Petersen et al., 2003; Hasenstaub et al., 2007). Furthermore, whiskers are known as tactile elements, used by rodents in social settings (Lenschow et al., 2015), for detecting the location and presence of objects (Hutson and Masterton, 1986), for discriminating between fine textures (Carvell and Simons,1995; Chen et al., 2013; Kerekes et al., 2017), and are used to guide mice as they navigate their environment (Hutson and Masterton, 1986; Carvell and Simons, 1990; Brecht et al., 1997; MItchinson et al., 2011; Grant et al., 2012; Voigts et al., 2015). In the plus maze used here, the floor and walls of the maze, their texture, the edges of the lane openings can all provide tactile input. Whiskers contact the wall as the animal navigates the maze, and this contact might result in the asymmetry. But our work shows that while whiskers do invariably contact the walls (Figure 1C, in left turn and forward motion and **Videos 1-3**), contact isn’t the main reason for side to side asymmetry; the side to side asymmetry persists even after whiskers have been trimmed to stubs that provide little tactile input. This suggests that the asymmetry arises as a postural adjustment or as a preparation for the actions that can follow hundreds of milliseconds after the initial asymmetric positioning of whiskers. That the level of asymmetry, and the setpoint / amplitude of whisking on two sides of the face, increases and decreases in a behaviorally relevant fashion, i.e. it increases as the animal approaches a turn, and the protracted and retracted side flip as the animal makes the turn, suggest that this asymmetry is related to active positioning of the whiskers (Figure 7).

The persistence of a correlation between asymmetric positioning and motion of whiskers and behavioral state even after trimming of the whiskers implies that tactile input from the walls does not drive the asymmetries, and tactile input does not drive the relationship to behavioral states. Whisking in rodents has been synonymous with exploration, navigation, and the way whiskers spread around the face during active tactile exploration has been described as forming a canopy of mobile sensors (Welker, 1964; Wineski, 1983; Carvell and Simons, 1990). Whisking has also been linked to the expression of overt attention to points in space around the face and head (Mitchinson and Prescott, 2013), and whisking strategy has been linked to the novelty of the environment (Arkley et al., 2014).

The prevalence of whisking asymmetry in the Airtrack is striking: it is evident just as mice begin to move the maze and it is evident throughout their behavior in the maze. Previous work in head fixed mice has shown that whisking asymmetry relates to the direction of movement of the animal on an air ball (Sofronview et al., 2014). In contrast in freely moving mice asymmetry relates to multiple factors: anticipation of future head movements (Towal and Hartmann, 2006); anticipation of contact (Grant et al., 209; Mitchinson et al, 2011; Voigts et al., 2015) and contact (Mitchinson et al., 2013; Schroeder and Ritt, 2016). Our results fit with anticipatory asymmetry that has nothing to with head movement, or whisker contact, but is related to the direction of movement of the maze and potentially related to the direction of the movement of the mouse.

It is possible that motor preparation involves changes in breathing triggered by anticipation of the additional effort that walking involves, and this change in breathing triggers nose movement, and whisking that co-occurs with each bout of movement the animal makes (Kurnikava et al., 2017). But asymmetry in whisker set point probably does not arise from breathing or sniffing which are fast events and occur simultaneously in both nares.

In sum, our work suggests that it is possible to discern the internal behavioral state of the animal from the whisker position (Figure 7) and that brain circuits that control the animal’s motion, whisking and motion of the nose are engaged together but distinctly during each behavioral epoch. The changes in whisking reflect changes in the state of the animal, with the set point and whisking amplitude independently reflecting internal brain states. In human beings, facial movements are used for communication of emotion: distress, pleasure anger. Whether facial expression in mice reflects emotions, attention, or another aspect of behavior is only beginning to be explored. The degree of correlation between whisker position and the animals behavioral state reveals how whisking is interweaved with a number of other behaviors including breathing, and sniffing and head movement. Astonishingly whisking asymmetry occurs spontaneously, it is not instructed, and is probably related to activity in widespread areas of cortex (Stringer et al, 2019; Musall 2018). It is therefore even more remarkable that the internal state of the animal is predicted by the positioning of, and motion of these tactile sensory organs.

## Supporting information

Whisker tracking in task

Behavioral epochs and whisker movement

Freely moving mouse 2

Wide angle view of platform

Freely moving mouse 1

## Acknowledgements

We thank the Charité Workshop for technical assistance especially Alexander Schill, Jan-Erik Ode and Daniel Deblitz. We also thank members of the Larkum lab, specially Naoya Takahashi and Jaan Aru, for useful discussions about earlier versions of this manuscript.

## Videos

**Video 1.** A wide-angle view of a mouse performing a single trial in the maze. In this trial, the mouse backs out of a lane, and partially enters a lane where the LED light is still on, then backs out and enters a dark lane and gets rewarded for correct performance. The nose and two whiskers were painted bilaterally.

**Video 2.** A close up view of a mouse performing a single trial. In this trial, the mouse backs out of a lane, and enters an incorrect lane and waits there, then backs out and enters a dark lane and gets rewarded for correct performance.

**Video 3**. Video of the masks used for each whisker and nose in a single trial.

**Video 4**. Top view of whisker positions in each distinct behavioral epoch.

**Videos 5-6.** Freely moving mice. Whiksers were tracked in mice that had been trained in the maze while head fixed. Whisker positions can be seen as mice move forward and backward. Note that the top of the maze is covered with clear plexiglass.

**Supplementary Figure 1.**
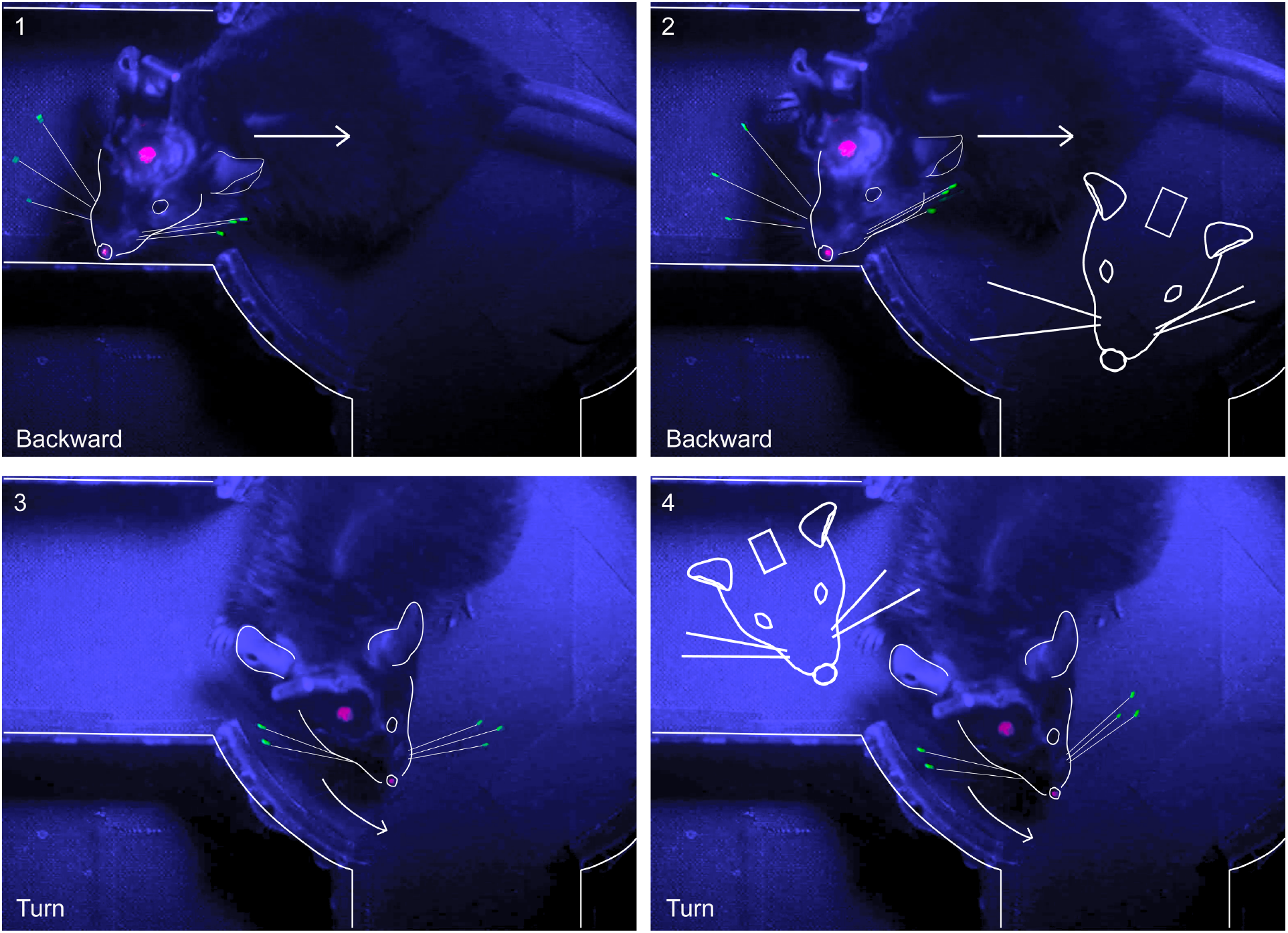
Asymmetry in freely moving animals. Mice that had previously been trained in the Air-Track were imaged navigating the maze without head-fixation. While whisker asymmetry persisted in freely moving animals and looked qualitatively similar to that seen in head fixed mice, it was also associated with changes in head position. The head and whiskers have been outlined here to make it easier to see whisker position and the animals head rotation. The direction that the animal was moving in is given by the arrows. In freely moving animals, head rotation, and angle relative to body also played a role, but the side to side asymmetry was still pronounced and similar to that seen in animals that were head fixed. The white schematic of the mouse (two right panels) shows the position of whiskers in a head-fixed animal at the same location in maze.

